# Duplication and divergence of Sox genes in spiders

**DOI:** 10.1101/212647

**Authors:** Christian L. B. Paese, Daniel J. Leite, Anna Schoenauer, Alistair P. McGregor, Steven Russell

**Affiliations:** Department of Biological and Medical Sciences, Oxford Brookes University, Gipsy Lane, Oxford, OX3 0BP, UK; Department of Genetics, University of Cambridge, Downing Street, Cambridge, CB2 3EH, UK

**Keywords:** Sox genes, *Parasteatoda tepidariorum*, *Stegodyphus mimosarum* spider, evolution, development

## Abstract

**Background:** The Sox family of transcription factors are present and conserved in the genomes of all metazoans examined to data and are known to play important developmental roles in vertebrates and insects. However, outside the commonly studied *Drosophila* model little is known about the extent or conservation of the Sox family in other arthropod species. Here we characterise the Sox family in two chelicerate species, the spiders *Parasteatoda tepidariorum* and *Stegodyphus mimosarum*, which have experienced a whole genome duplication (WGD) in their evolutionary history.

**Results:** We find that virtually all of the duplicate Sox genes have been retained in these spiders after the WGD. Analysis of the expression of Sox genes in *P. tepidariorum* embryos indicates that it is likely that some of these genes have neofunctionalised after duplication. Our expression analysis also strengthens the view that an orthologue of vertebrate Group B1 genes, *SoxNeuro*, is implicated in the earliest events of CNS specification in both vertebrates and invertebrates. In addition, a gene in the *Dichaete/Sox21b* class is dynamically expressed in the spider segment addition zone, suggestive of an ancient regulatory mechanism controlling arthropod segmentation as recently suggested for flies and beetles. Together with the recent analysis of Sox gene expression in the embryos of other arthropods, our findings are also indicative of conserved functions for some of these genes, including a role for *SoxC* and *SoxD* genes in CNS development, *SoxF* in limb development and a tantalising suggestion that *SoxE* genes may be involved in gonadogenesis across the metazoa.

*Conclusions:* Our study provides a new chelicerate perspective to understanding the evolution and function of Sox genes and how the retention of duplicates of such important tool-box genes after WGD has contributed to different aspects of spider embryogenesis. Future characterisation of the function of these genes in spiders will help us to better understand the evolution of the regulation of important developmental processes in arthropods and other metazoans including neurogenesis and segmentation.

## Introduction

The evolution of metazoan life forms was in part driven by the acquisition of novel families of transcription factors and signalling molecules that were subsequently expanded by gene duplications and evolved new functions [1, 2]. One such family, encoded by Sox genes, encompasses a set of conserved metazoan specific transcriptional regulators that play critical roles in a range of important developmental processes [3], in particular, aspects of stem cell biology and nervous system development [4, 5].

The Sox family is defined by a set of genes containing an HMG class DNA binding domain sharing greater than 50% sequence identity with that of SRY, the Y-linked sex determining factor in eutherian mammals [6]. In the chordates the family contains approximately 20 genes, which have been subdivided into eight groups (A-H) based mainly on homology within the DNA binding domain but also related group-specific domains outwith the HMG domain [7, 8]. In all metazoans examined to date representatives of the Sox family have been identified and these are largely restricted to Groups B to F [9] with other groups specific to particular lineages. While Sox-like sequences have been reported in the genome of the choanoflagellate *Monosiga brevicollis* [10] these are more closely related to the non-sequence specific HMG1/2 class of DNA binding domain and thus true Sox genes are restricted to metazoans [11, 12].

While vertebrate Sox genes have been intensively studied due to their critical roles in development [3], with the exception of the fruit fly *Drosophila melanogaster*, they are less well characterised in invertebrates. *D. melanogaster* contains eight Sox genes (four group B and one each in groups C to F), which is generally consistent across the insect genomes examined to date [9, 13, 14]. Of particular interest are the Group B genes of insects, which share a common genomic organisation that has been conserved across all insects examined to date, with three genes closely linked in a cluster [13–15]. Critical roles in early segmentation and nervous system development have been shown for *Dichaete* (*D*) [16, 17]}, and in CNS development for *SoxNeuro* (*SoxN*), where both these group B genes show partial redundancy [18, 19]. The evolutionary conservation of Sox protein function as well as sequence has been shown in rescue or swap experiments, where mouse Sox2 rescues *D* null mutant phenotypes in the *D. melanogaster* embryo [20] and *Drosophila* SoxN can replace Sox2 in mouse ES cells [21]. Furthermore, a comparison of D and SoxN genomic binding in the *D. melanogaster* embryo with Sox2 and Sox3 binding in mouse embryonic or neural stem cells indicates that these proteins share a common set of over 2000 core target genes [22–24]. These and other studies suggest that Sox proteins have ancient roles, particularly in the CNS, where their functions have been conserved from flies to mammals.

Of the other two *D. melanogaster* group B genes, *Sox21a* plays a repressive role in maintaining adult intestinal stem cell populations [25, 26] but there is no known function for *Sox21b.* The group C gene *Sox14* is involved in the response to the steroid hormone ecdysone and is necessary for metamorphosis [27]; *Sox102F* (Group D) has a role in late neuronal differentiation [28]; *Sox100B* (Group E) is involved in male testis development [29] and *Sox15* (Group F) is involved in wing metamorphosis and adult sensory organ development [30, 31].

While functional studies are lacking in other insects, gene expression analysis in *Apis mellifera* and *Bombyx mori* indicates that aspects of Sox function are likely to be conserved across species [13, 14]. More recently, a conserved role for *D* in the early embryonic segmentation of both *Drosophila* and the flour beetle *Tribolium castaneum* suggests that aspects of regulatory function as well as genomic organisation may have been conserved across insects [32]. Outside the insects little is known, however genome sequence analysis and gene expression studies suggest key roles for Sox family members in stem cell and cell fate processes in Ctenophores [12] and Porifera [33], as well as neural progenitor development in Cnidarians [34] and a Dioplopod [35]. Taken together with the extensive work in vertebrate systems, it is clear that Sox genes play critical roles in many aspects of metazoan development, at least some of which appear to be deeply conserved.

Arthropods comprise approximately 80% of living animal species [36] exhibiting a huge range of biological and morphological diversity that is believed to have originated during the Cambrian Period over 500 million years ago [37]. While the analysis of traditional model arthropods such as *D. melanogaster* has taught us much about conserved developmental genes and processes, it is only more recently that genomic and other experimental approaches are beginning to shed light on the way genes and regulatory networks are deployed to generate the diversity of body plans found in other insects [38] and more widely in chelicerates and myriapods [39]. In terms of the Sox family, recent work indicates conserved Group B expression in the early neuroectoderm of the myriapod *Glomeris marginata* [35] and neurectodermal expression of a Group B gene in the chelicerate *P. tepidariorum* has been reported [40].

Chelicerates in particular offer an interesting system for exploring the evolution and diversification of developmental genes since it has emerged that some arachnid lineages, including spiders and scorpions, have undergone a whole genome duplication (WGD) [41]. Interestingly, duplicated copies of many developmental genes, including Hox genes and other regulatory factors such as microRNAs, have been retained in *P. tepidariorum* and other arachnids [41, 42]. Thus, chelicerate genomes provide an opportunity to explore issues of gene retention, loss or diversification [43].

Here we report an analysis of the Sox gene family in the spiders, *P. tepidariorum* and *S. mimosarum*, and show that most duplicate Sox genes have been retained in the genomes of these spiders after the WGD. Furthermore, while group B genes show highly conserved expression in the developing CNS, the expression of other spider Sox genes suggests they play important roles and potentially novel functions in other aspects of embryogenesis.

## Results and Discussion

### Characterisation of Sox genes in spiders

In order to characterise the Sox gene complement of spiders we conducted TBLASTN searches of the genomes of *P. tepidariorum* [41] and *S. mimosarum* [44] using the HMG domain of the mouse Sox2 protein, recovering 16 and 15 sequences respectively. All but three of these contained the highly conserved RPMNAFMVW motif that is characteristic of Sox proteins and these three (*ptSoxC-2*, *ptSoxB-like* and *ptSox21b-2*) only show minor conservative substitutions in this motif. 14 of the *P. tepidariorum* sequences corresponded with annotated gene models. Two sequences were identical (*ptSox21b-1*, aug3.24914.t1 and aug3.g24896.t1), since the latter maps to a genomic scaffold of only ~7 kb we presume this represents an assembly error and thus consider them as a single gene. One genomic scaffold encoding a Sox domain (*ptSoxB-*like, Scaffold3643:28071..28299) is in a region of poor sequence quality and we cannot be sure it represents a *bona fide* gene but have nevertheless included it in subsequent analysis. In the case of *S. mimosarum* we identified 15 genomic regions, 11 of which correspond to annotated genes. Reciprocal BLAST searches of *D. melanogaster* or vertebrate genes recovered Sox proteins as top scoring hits. In addition to these true Sox gene sequences, we also recovered sequences that correspond to the *D. melanogaster capicua* (*cic*) and *bobby sox* (*bbx*) genes but do not consider these Sox-related genes further here.

To classify the spider Sox proteins we generated MUSCLE sequence alignments and PhyML maximum likelihood phylogenies using the HMG domains recovered from the BLAST searches, along with those from the eight *D. melanogaster* Sox genes and representatives of each subgroup from mouse (Supplementary Table 1). These analyses resulted in a clear classification into groups B-F as found in other invertebrate genomes (Figure 1). Note that Group A only contains the *SRY* gene specific to eutherian mammals and there are no Group G, H or I Sox genes found outside the vertebrates. Supporting this classification, phylogenetic trees constructed with the full length sequences of the predicted spider Sox proteins and those from *D. melanogaster* yielded virtually identical results (Supplementary Figure 1). Following the recommended nomenclature for Sox genes [7], we have named the spider Sox genes as indicated in Supplementary Table 1. The naming of *D. melanogaster* Sox genes is confusing with some carrying historic names based on phenotype (*Dichaete* and *SoxN*), others named after cytological locations (*Sox100B* and *Sox102F*) and others with inappropriate numerical designations (*D. melanogaster Sox14* is a Group C gene while the vertebrate Sox14s are in Group B and *D. melanogaster Sox15* is in group F, while vertebrate Sox15s are in Group G), thus we propose renaming the *D. melanogaster* group C-F genes according to the standard nomenclature used in the Sox field: these designations are already recognised as synonyms in FlyBase. With respect to the Group B genes, since the sequence and organisation of these appears to be invertebrate specific, we propose a nomenclature based on the current *D. melanogaster* gene names: *SoxN*, *D*, *Sox21a* and *Sox21b* (Supplementary Table 1).

**Figure 1.**
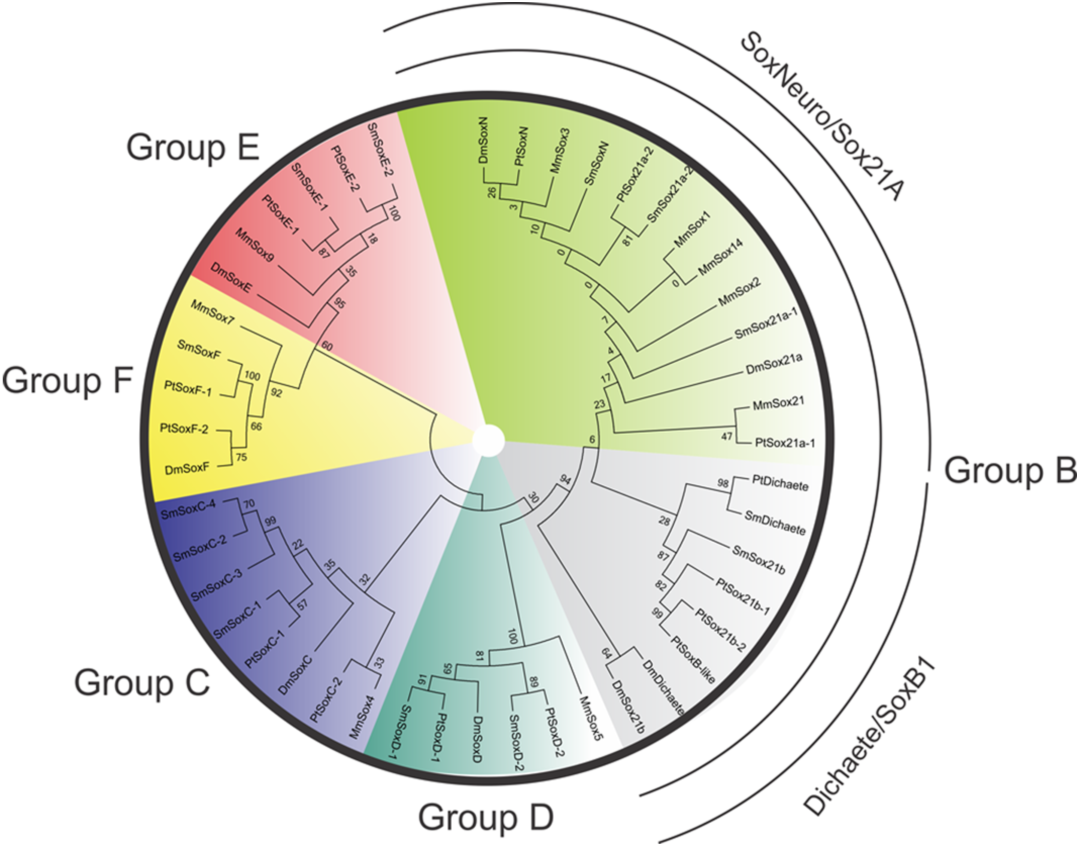
Phylogeny of Sox HMG domains in selected metazoans. Phylogenetic tree showing the relationship between *Mus musculus* (Mm), *D. melanogaster* (Dm), *P. tepidariorum* (Pt) and *S. mimosarum* (Sm) Sox genes based on HMG domain sequences. The grouped genes are divided into different colours as highlighted outside the circle.

In common with many other gene families in spiders [41], the Sox genes are mostly represented by two or more copies in each group (Figure 2). In other arthropods examined to date, as well as the onochophoran *Euperipatoides kanangrensis* [45], there is usually only a single copy of each gene, although [45] recently reported two Group E genes in the millipede *G. marginata*. In the case of spider Groups D and E, the duplications clearly predate the divergence of the two spider species we analysed since the duplicates group together in the phylogenetic analysis and show extensive homology across the length of the coding sequence (Figure 1). With Group F, there is only one gene identified in *S. mimosarum* but two in *P. tepidariorum*. In the case of group C, there appears to have been additional duplication events in *S. mimosarum*. When we consider the full-length protein sequences (Supplementary Figure 1), *ptSoxC-1* groups with *smSoxC-1* and *ptSoxC-2* with *smSoxC-2*. *smSoxC-2* has undergone a local head-to-head duplication, with *ptSoxC-2* and *smSoxC-3* adjacent in the genome. *smSoxC-4* has no predicted gene model but the region of the genome encodes an uninterrupted HMG domain closely related to those of the *smSoxC-2* and *C-3* duplicates. Whether this is a *bona fide* gene remains to be determined.

**Figure 2.**
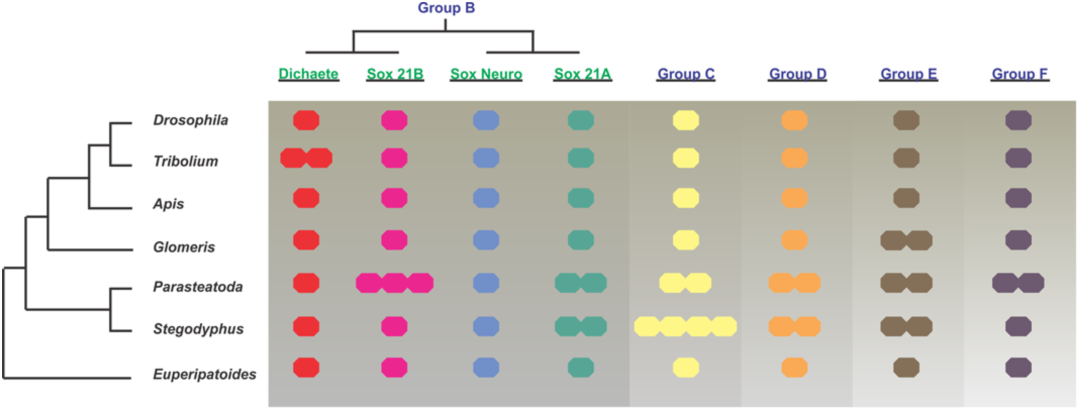
Repertoire of Sox genes in selected arthropods. Diagrammatic representation of the complement of Sox genes in insects (*Drosophila melanogaster*, *Tribolium castaneum* and *Apis mellifera*), the spiders (*Parasteatoda tepidariorum* and *Stegodyphus mimosarum*), the myriapod (*Glomeris marginata*) and an onychophoran (*Euperipatoides kanangrensis*). Each coloured circle represents a gene.

In many organisms, some genes in Groups D, E and F contain an intron within the DNA binding domain sequence in a position that is highly conserved and specific for each group [7]: our analysis indicates that this is also the case for the spider genes in these three groups (see arrows in Figure 3). Interestingly, while there is an intron within the spider Group D genes, it has been lost in the *D. melanogaster* orthologue. Secondary intron loss is also observed in Group F, where mouse *Sox7* has no intron but the related *Sox17* and *Sox18* genes do. The location of these HMG domain introns suggests they were present in the common ancestor of the vertebrates and the arthropods.

**Figure 3.**
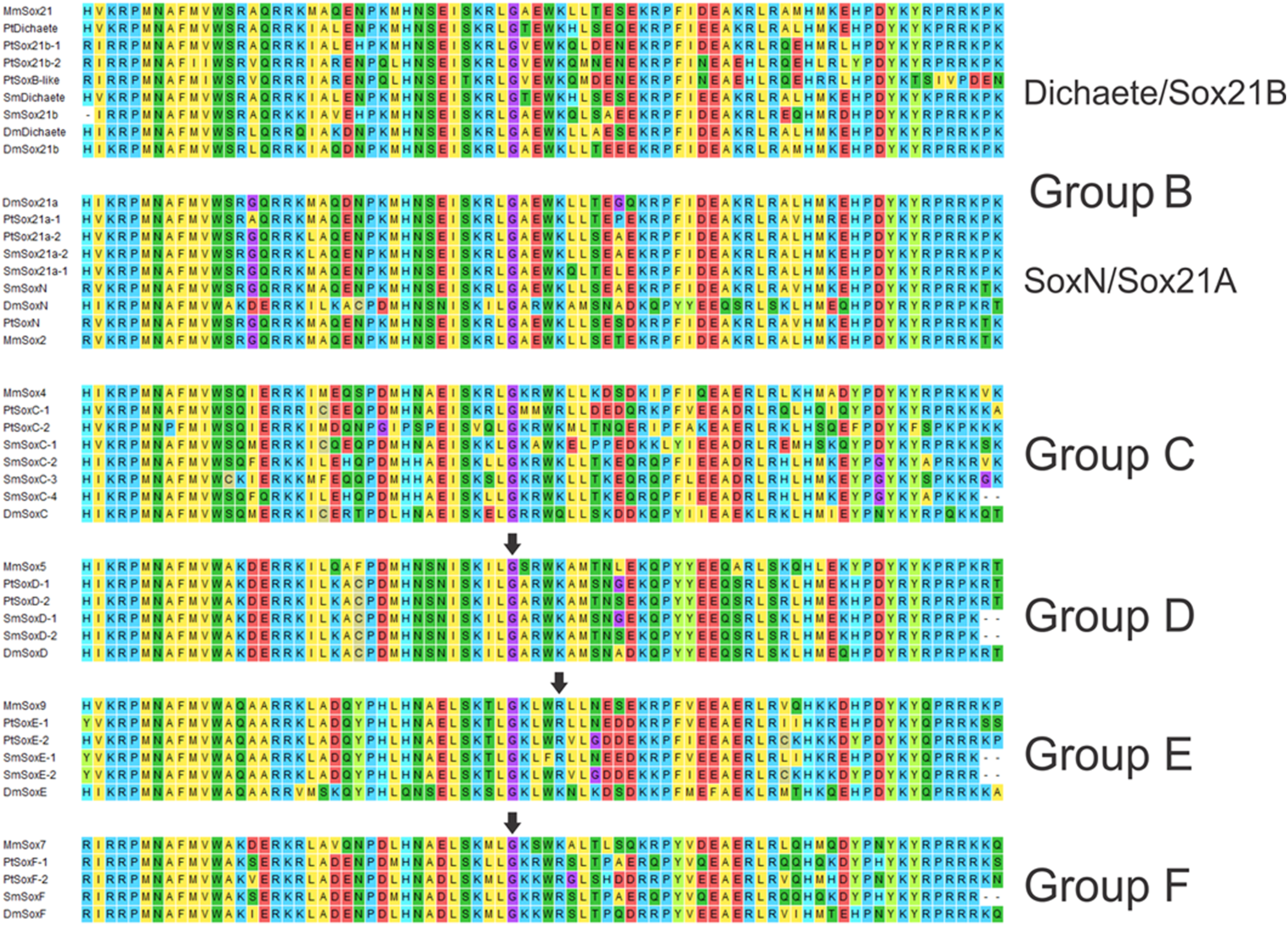
Multiple sequence alignment of HMG domain sequences from selected metazoans. *Mus musculus* (Mm), *D. melanogaster* (Dm), *P. tepidariorum* (Pt) and *S. mimosarum* (Sm). Arrowheads indicate the locations of HMG domain introns and the bold underlined amino acids indicate the genes where the introns are present.

The Group B genes of insects and vertebrates are clearly different. Vertebrate Group B genes are subdivided into B1 (Sox1, 2 and 3) and B2 (Sox14 and 21), a classification manifest both at the sequence and the functional levels, with Group B1 proteins acting as transcriptional activators particularly important for nervous system specification, while the Group B2 proteins act as transcriptional repressors [46–48]. In contrast, the organisation and functional classification of Group B genes in insects is subject to some debate. There is a clear orthologue of the Group B1 proteins, represented by *SoxN* in *D. melanogaster* and genes named *SoxB1* or *Sox2* in every invertebrate genome examined. The remaining three *D. melanogaster* Group B genes (*D*, *Sox21* and *Sox21b*) have been characterised as Group B2 based on sequence alignments with vertebrate proteins. In *D. melanogaster* these three genes are arranged in a cluster on Chromosome *3L*, an organisation that is conserved across at least 300 MY of evolution, with a similar organisation found in flies, mosquitoes, wasps, bees and beetles [11, 13, 15]. While there is evidence that *Sox21a* has a repressive role consistent with the vertebrate B2 class [25, 26], considerable genomic evidence clearly shows D acts as a transcriptional activator, a role inconsistent with that observed for vertebrate SoxB2 proteins [22, 49].

The phylogenies generated with the HMG domains from a range of species (Figure 1; Supplementary Figure 2) or full length proteins sequences from spiders and *D. melanogaster* (Supplementary Figure 1) support a classification of arthropod Group B genes where there is a single *SoxN* gene, one or more *Sox21a* genes and two or more *Dichaete-Sox21b* genes. In spiders we find strong support for a single *SoxN* gene, duplications of the *Sox21a* class and a single *D*-like gene in both species. In *P. tepidariorum* we find a duplication of the *Sox21b* genes and the possibility of a further tandem duplication of *ptSox21b-2* gene if the *ptSoxB-like* ORF is a genuine gene. *S. mimosarum*, in contrast has a single *Sox21b* class gene. Intriguingly, we find that two *P. tepidariorum* Group B genes (*ptDichaete* and *ptSox21a-*1) are located in the same genomic region, separated by over 200 kb of intervening DNA that is devoid of other predicted genes (Figure 4), an organisation reminiscent of that found in insects. Indeed, the linkage of *ptDichaete* and *ptSox21a-1* supports the idea that these genes were formed by a tandem duplication in the protostome/deuterostome ancestor [11, 15]. The separation of *SoxN* from the *D/Sox21a-1* cluster in the spider suggests that either this fragmentation happened early in arthropod evolution [11] or that the duplication and separation of *SoxN* and *D* (or *Sox21a*) occurred early in Sox evolution [11, 15] (Figure 4).

**Figure 4.**
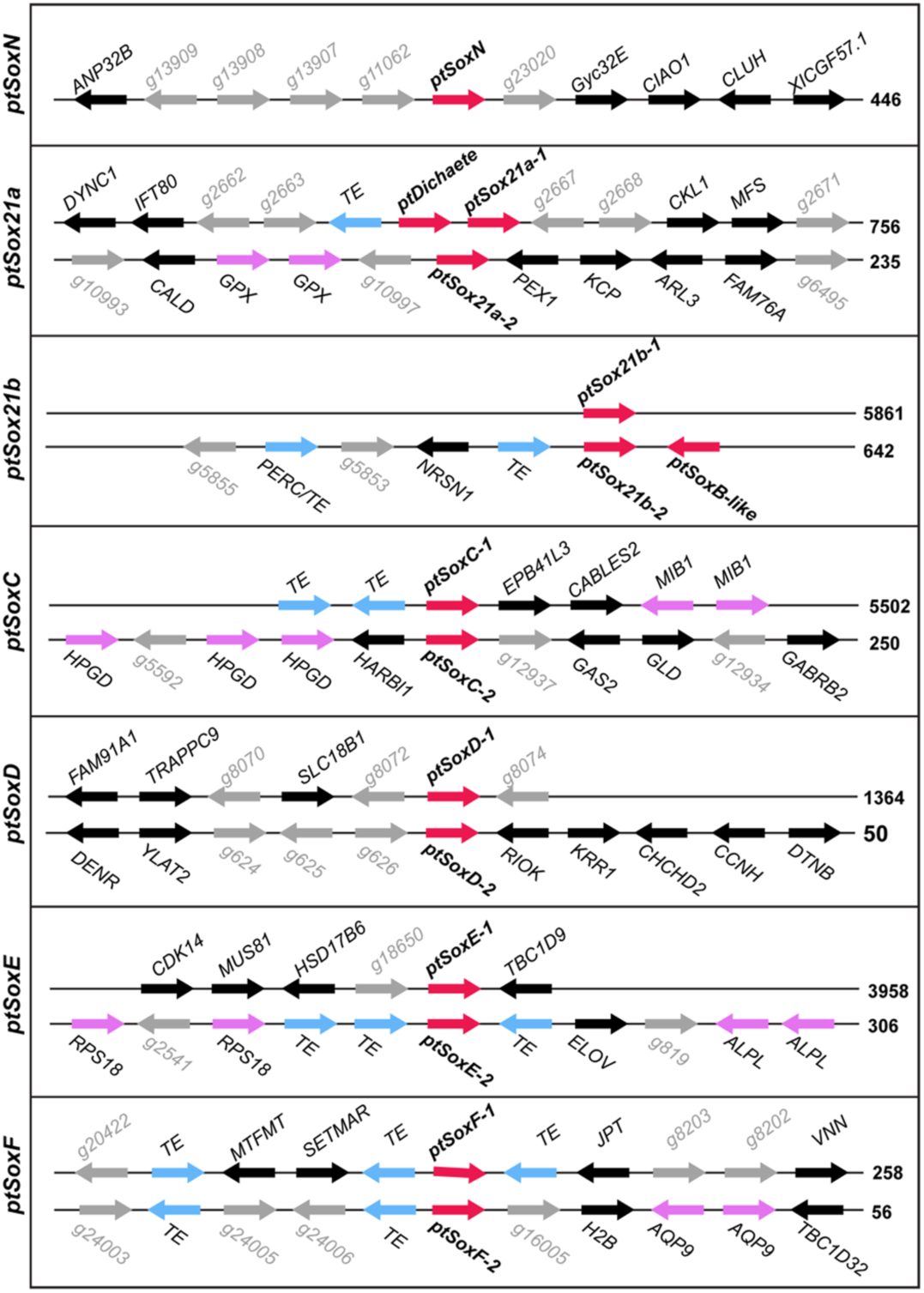
Sox gene synteny in the *P. tepidariorum* genome. The synteny of Sox genes (red) and flanking genes that have putative homology (black) compared between the Sox paralogs. Homology of flanking genes was also used to indicate tandem duplicates (pink), transposable elements (TEs) (blue). Genes that lack homology are shown in grey with their gene model IDs. Only the SoxF genes were found in the same transcriptional orientation as upstream TEs. Of the thirteen Sox containing scaffolds, six scaffolds contained TEs that flank the Sox genes. Transcriptional direction is indicated by arrows. The DoveTail/HiRise scaffold ID numbers are given on the right.

Taken together, our analysis clearly shows that the spider genomes we examined have the full complement of Sox genes found in insects, mostly retained duplicates in Groups C, D, E and F after the WGD, and have a Group B organisation that more closely resembles insects than vertebrates.

### Arrangement of P. tepidariorum Sox genes after WGD

The phylogenetic relationships of Sox genes in *P. tepidariorum* suggest that there are two paralogs of each Sox group except for *SoxN* and *D* (Figures 1 and 2). To investigate if all of these duplicated Sox genes arose from the WGD event in the ancestor of these animals [41], the synteny of Sox genes was analysed in the *P. tepidariorum* genome (Figure 4).

Most of the Sox genes in *P. tepidariorum* were found dispersed in the genome on separate scaffolds consistent with the expectation that they arose via WGD. Analysis of the five upstream and five downstream genes flanking each Sox gene, however, revealed that dispersed duplicated Sox genes are generally not closely linked to other duplicated genes (Figure 4, Supplementary Table 2). While it is likely that this is a consequence of extensive loss of ohnologs and genomic rearrangements since the WGD 430 MYA, we cannot rule out that at least some of the duplicated Sox genes in this spider arose via tandem duplication followed by rearrangements after the WGD. The only tentative example of retained synteny was in the SoxF group where we found that the two *SoxF* genes of *p. tepidariorum* have an upstream flanking sequence with homology to a transposable element (TE) with matching transcriptional orientation. Interestingly, six of the thirteen Sox containing scaffolds also have TE-like sequences nearby (Figure 4). TEs have previously been linked to the expansion of genes and their rearrangements [50, 51], however further analysis is needed to determine if TEs identified in this synteny analysis are involved in the evolution of Sox genes in spiders.

The exceptions to the dispersion of Sox genes in *P. tepidariorum* are *ptDichaete* and *ptSox21a-1* on scaffold #756 (as discussed above), as well as *ptSox21b-2* and *SoxB-like* on scaffold #642 (Figure 4). The sequences of the HMG domains of the clustered *ptSox21b-2* and *SoxB-like* genes grouped together with high bootstrap confidence indicative of a head-to-head tandem duplication (Figures 1 and 4). However, the HMG domain of *SoxB-like* is split across two reading frames and although the sequence quality is poor in parts of this scaffold, it's sequence similarity to *ptSox21b-2* suggests that *SoxB-like* may have been pseudogenised (Figure 4).

### Sox Gene Expression during P. tepidariorum embryogenesis

We next studied the expression of Sox genes during embryogenesis in *P. tepidariorum.* For the SoxB family genes *ptSox21a-1*, *ptSox21a-2*, *ptSox21b-2* and *D*, we did not detect any expression during embryogenesis. This might indicate that they are only expressed at very low levels or in a few cells or that these genes are used during post-embryonic development.

*ptSoxN* expression is visible from late stage 7 in the most anterior part of the germ band, a region corresponding to the presumptive neuroectoderm (Figure 5A). This head-specific expression in *P. tepidariorum* is similar to early expression of *SoxN* observed in *D. melanogaster* [52] and in *A. mellifera*, where *SoxB1* is expressed in the gastrulation fold and the anterior part of the presumptive neuroectoderm [13]. *ptSoxN* is subsequently expressed broadly in the developing head and follows neurogenesis in a progressive anterior-to-posterior pattern as new segments are added (Figure 5B). By mid stage 9, *ptSoxN* is strongly expressed in the head lobes and in the ventral nerve cord (Figure 5C), however, after this stage no further expression was detected. In both *D. melanogaster* and *A. mellifera, SoxN* expression is also observed throughout the neuroectoderm and becomes restricted to the neuroblasts [13, 18, 19].

**Figure 5.**
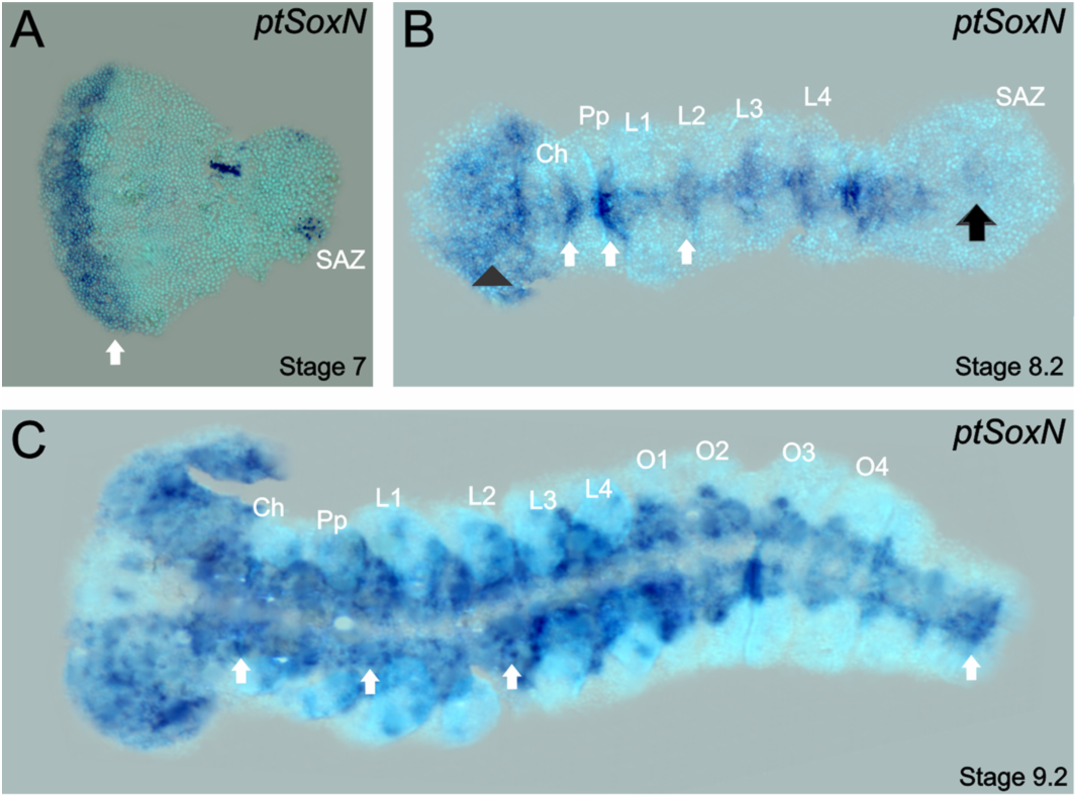
Expression of *ptSoxN*. Flat-mount embryos at different stages of development after RNA *in situ* hybridization. A) *ptSoxN* expression is restricted to the presumptive neuroectoderm in the most anterior region of the germ band in stage 7 embryos (arrow). B) At stage 8.2, expression is in the most anterior part of the embryo (black arrowhead) and in the ventral nerve cord appearing sequentially from anterior to posterior, white arrows indicate expression in clusters that will subsequently broaden, expression in the posterior region adjacent to the SAZ is also observed (black arrowhead). C) At stage 9.2 expression is observed throughout the ventral nerve cord, with differentiating clusters indicated by arrows. Ch: chelicerae, L1 – L4: prosomal segments 1 to 4, O1 – O4: opisthosomal segments 1 to 4, Pp: pedipalps; SAZ: segment addition zone. Ventral views are shown for all embryos with the anterior to the left.

In chelicerates, neurogenic progenitors were shown to delaminate in clusters of cells rather than single neuroblast-like cells found in dipterans and some hymenopterans [53]. However, even with these different modes of neurogenic differentiation, the expression of *SoxN* orthologues suggests this gene performs the same function. Indeed, the recent study by [45], of *T. castaneum*, *E. kanangrensis* and *G. marginata* also shows that the *SoxN* orthologues in these species have widespread and early neuroectodermal expression. Taken together these data clearly support the view that throughout the Bilateria a SoxN class protein is a marker of the earliest stages of neural specification.

Another member of the B group, *ptSox21b-1,* shows dynamic expression in the nascent prosomal segments and in the posterior segment addition zone (SAZ) from stage 7 (Figure 6A and B). At stage 8.2 expression is observed in the most anterior part of the germ band, which corresponds to the presumptive neuroectoderm in the future head and prosomal segments (Figure 6C). At stages 9 and 10, strong expression is apparent throughout the ventral nerve cord, similar to *ptSoxN*, and cyclical expression is also detected in the SAZ (Figure 6D and E).

**Figure 6.**
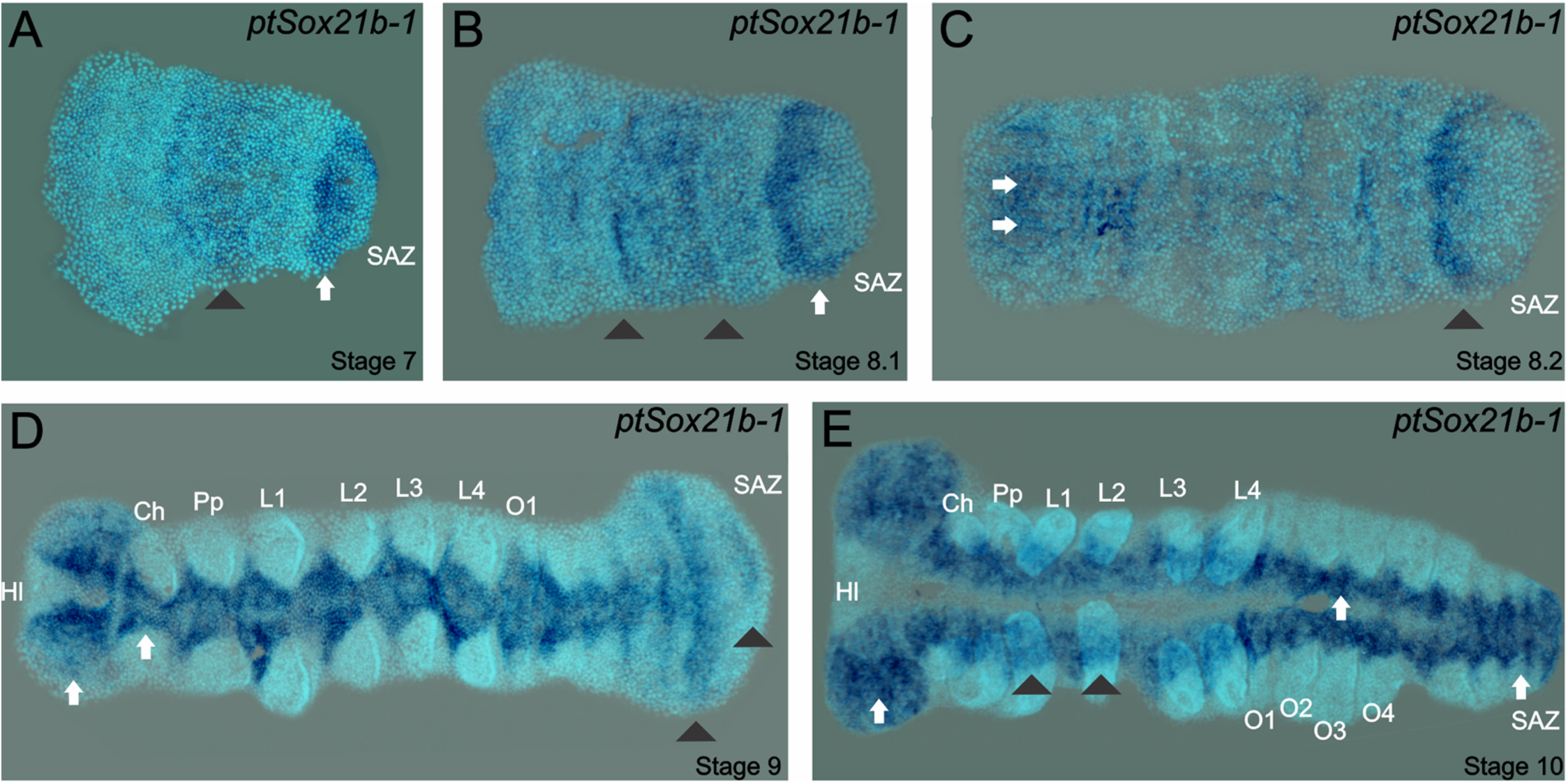
Expression of *ptSox21b.* Flat-mount embryos at different stages of development after RNA *in situ* hybridization. A) *ptSox21b-1* expression is detected from mid-stage 7, where dynamic expression in the nascent segment (black arrowhead) and in the SAZ are indicated (white arrow). B) At stage 8.1, expression in the SAZ is dynamic (white arrow), and broadens in forming segments (black arrowheads). C) At stage 8.2, white arrows at the anterior indicate expression in the presumptive ventral nerve cord, with strong expression in the posterior SAZ still prominent (black arrowhead). D) At stage 9 strong expression in the entire anterior part of the ventral nerve cord is indicated by white arrows, expression is lower at the most posterior but remains dynamic in the SAZ (black arrowhead). E) At stage 10 expression is visible in the ventral nerve cord underneath the growing limb buds (black arrowheads) and becomes strong in the entire ventral nerve cord (white arrows). Ch: chelicerae, HL: head lobes, L1 – L4: prosomal segments 1 to 4, O1 – O4: opisthosomal segments 1 to 4, Pp: pedipalps; SAZ: segment addition zone. Ventral views are shown for all embryos with the anterior to the left.

In *T. castaneum*, *Sox21b* has similar expression to *D*, early in the SAZ and then in the developing CNS. In *E. kanangrensis* and *G. marginata,* there is no early *Sox21b* expression [45], however in these species, *D* is expressed during segmentation and then later in the CNS. This suggests that the role of *D* in segmentation in *D. melanogaster* and *T. castaneum* [32] could extend to *E. kanangrensis* and *G. marginata* but in spiders the closely related *Sox21b-1* gene may play this role.

For the Sox C group genes we did not detect any expression for *ptSoxC-2*. However, *ptSoxC-1* expression was detected at mid-stage 6, in a pattern similar to that of *ptSoxN* in the presumptive head and anterior segments (Figure 7A). By stage 8.2 expression is apparent in neuroectodermal progenitors along the germ band and at the anterior region of the SAZ (Figure 7B), however by stage 9.1 (Figure 7C) expression is lost from the SAZ. Interestingly, from stage 9.1, *ptSoxC-1* is expressed in the ventral nerve cord, from the head to the SAZ, however unlike the uniform expression of *ptSoxN*, *ptSoxC-1* is observed in clusters of cells, presumably undergoing neurogenic differentiation, progressively from the head through to opisthosomal segments as they differentiate in an anterior to posterior manner (Figure 7C).

**Figure 7.**
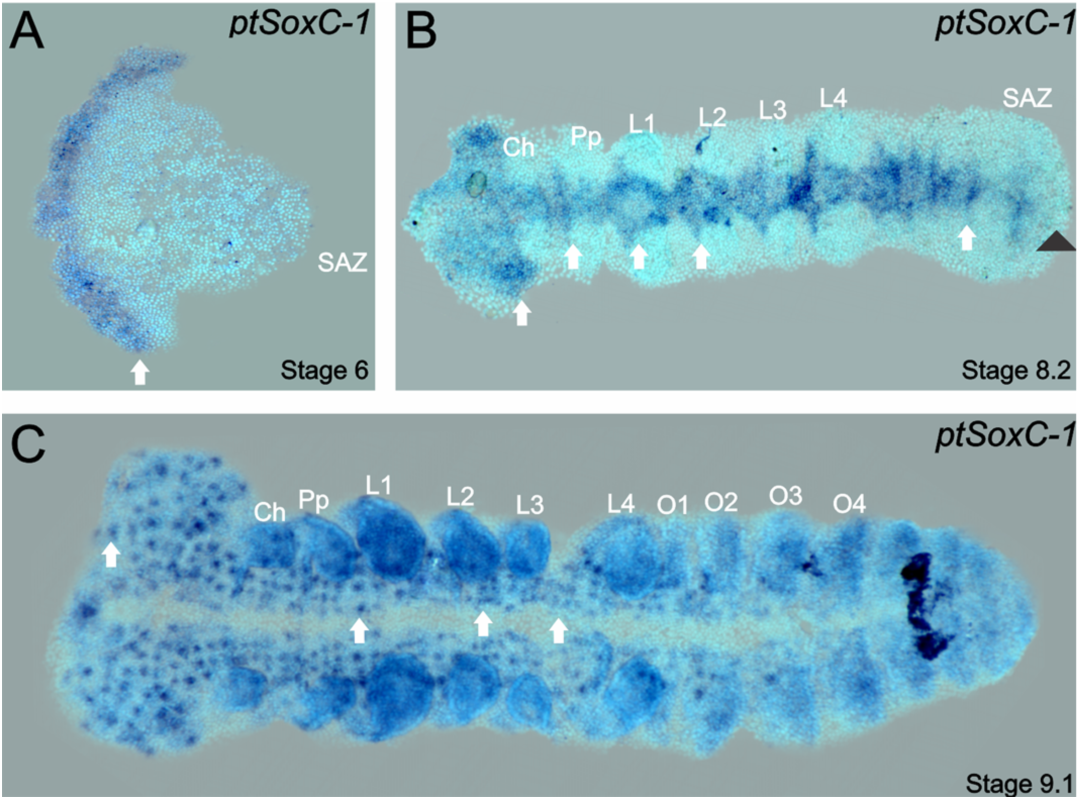
Expression of *ptSoxC-1.* Flat-mount embryos at different stages of development after RNA *in situ* hybridization. A) *ptSoxC-1* is strongly expressed in the presumptive neuroectoderm at stage 6 as indicated by the white arrow. B) At stage 8.2, strong expression is observed in the ventral nerve cord (white arrows) with the exception of the most posterior part of the SAZ (black arrowhead) C) At stage 9.1, expression is apparent in clusters of cells in the head and each anterior segment until the third opisthosomal segment (O3): white arrows indicate localized expression. The signal in the limb buds is background and staining at the most posterior part of the O5 segment is an artefact of incomplete chorion removal. Ch: chelicerae, L1 – L4: prosomal segments 1 to 4, O1 – O4: opisthosomal segments 1 to 4, Pp: pedipalps; SAZ: segment addition zone. Ventral views are shown for all embryos with the anterior to the left.

In *D. melanogaster* the single *SoxC* gene has been shown to play a role in the response to ecdysone at the onset of metamorphosis and has no known role in the embryonic CNS [27]. In contrast, the vertebrates *SoxC* genes (*Sox4, 11* and *12*) play critical roles in the differentiation of post-mitotic neurons, acting after the Group B genes, which specify neural progenitors [54]. In *A. mellifera,* late expression of the SoxC gene was observed in the embryonic cephalic lobes and in the mushroom bodies [13]. The expression of SoxC orthologues in the embryonic CNS of other invertebrates [45] suggests that this class of Sox gene may play a conserved role in aspects of neuronal differentiation, which has been lost in *D. melanogaster*. Interestingly, a comparison of target genes bound by Sox11 in differentiating mouse neurons and SoxN in the *D. melanogaster* embryo shows a conserved set of neural differentiation genes, suggesting that in *D. melanogaster* the role of *SoxC* in neuronogenesis has been taken over by *SoxN* [55].

We identified two genes in each of the SoxD, E and F families, however, we found no *in situ* evidence for expression of Sox*D-2, SoxE-2* or *SoxF-1* during the *P. tepidariorum* embryonic stages we examined. For *ptSoxD-2* we found no expression prior to stage 10, but we then observed expression in the ventral nerve cord from the head to the most posterior part of the opisthosoma (Figure 8A). The *D. melanogaster SoxD* gene is also expressed at later stages of embryonic CNS development [56] and has been shown to play roles in neurogenesis in the larval CNS [28]. While *SoxD* has been reported to be ubiquitously expressed in *A. mellifera* embryos, it is also expressed in the mushroom bodies of the adult brain [13]. Embryonic brain expression of SoxD orthologues in beetles, myriapods and velvet worms [45], as well as a known role for *SoxD* genes in aspects of vertebrate neurogenesis [54, 57] again suggests conserved roles for *SoxD* during metazoan evolution.

**Figure 8.**
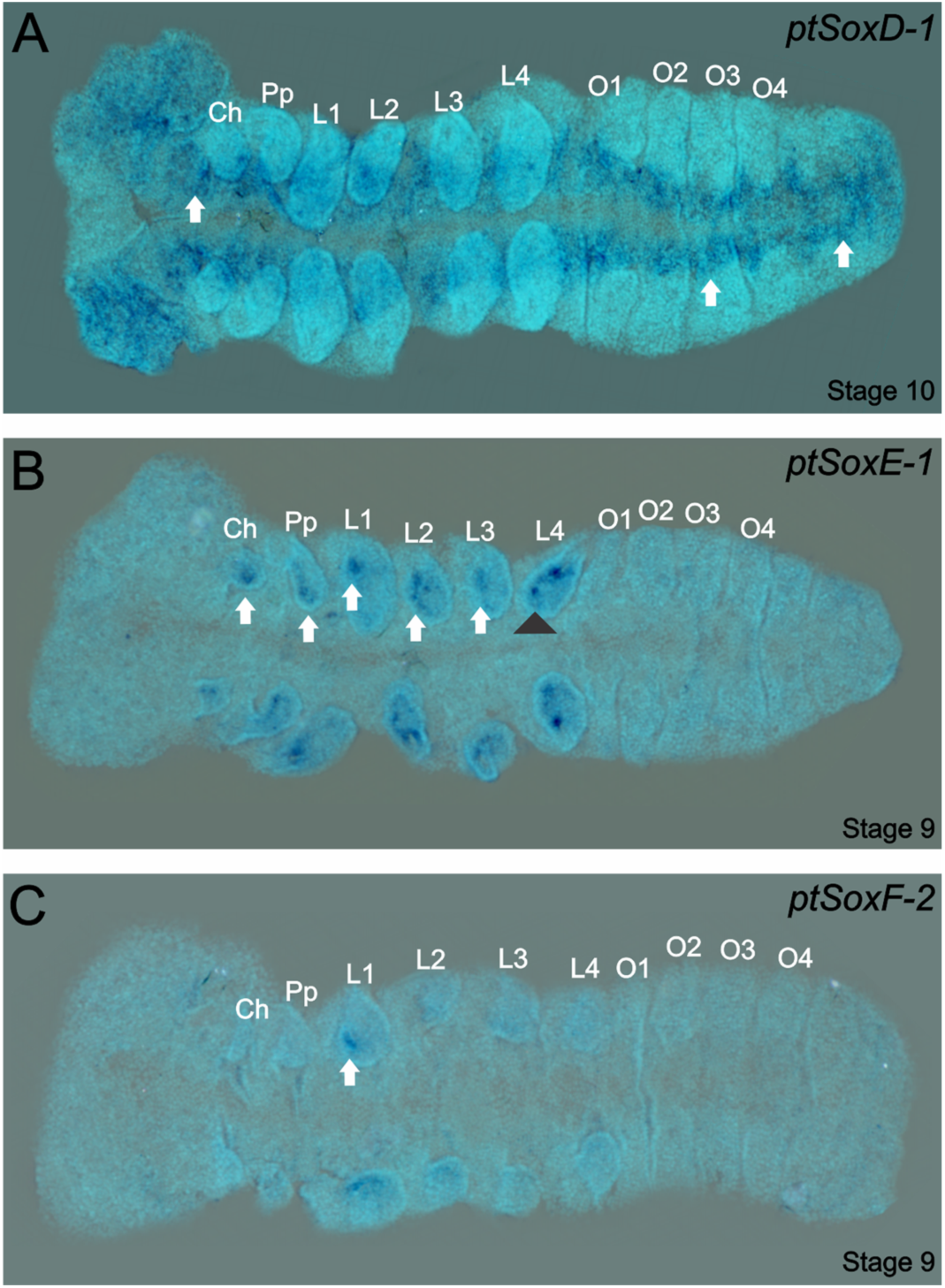
Expression of Sox D, E and F group orthologues. Flat-mount embryos at different stages of development after RNA *in situ* hybridization. A) *ptSoxD-1* expression is observed throughout the ventral nerve cord in stage 10 embryos as indicated by the arrows. B) *ptSoxE-1* expression at stage 9 is visible as single dots in the forming chelicerae, broader expression in the pedipalps and L1 to L3 (white arrows), and as two dots in the L4 limb bud as indicated by the black arrowhead. C) The expression of *ptSoxF-2* is only visible in the L1 limb buds forming at stage 9 (arrows). Ch: chelicerae, L1 – L4: prosomal segments 1 to 4, O1 – O4: opisthosomal segments 1 to 4, Pp: pedipalps; SAZ: segment addition zone. Ventral views are shown for all embryos with the anterior to the left.

*ptSoxE-1* is expressed in the developing limbs from stage 9 in small dots in the chelicerae, pedipalps and L1 buds, broader expression in L2 and L3, and in two dots in the L4 limb pairs, corresponding to the differentiating peripheral nervous system (PNS) (Figure 8B). We did not observed any expression of *ptSoxE-1* in opisthosomal segments 2 to 6 where the germline is believed to originate [58].

In *D. melanogaster* the *SoxE* orthologue is associated with both endodermal and mesodermal differentiation, is expressed in the embryonic gut, malpighian tubules and gonad [59], and has been shown to be required for testis differentiation during metamorphosis [29]. Both the *A. mellifera SoxE* genes are also expressed in the testis [13]. Janssen and colleagues [45], observed expression of *SoxE* genes in other invertebrates, associated with limb buds like in the spider, but they also detected posterior expression associated with gonadogenesis. These observations are particularly intriguing since the vertebrate *Sox9* gene has a crucial function in testis development [60]. Therefore, while we did not observe *SoxE* expression associated with early gonadogenesis it remains possible that the spider genes are used later in this process. We note that while the fly *SoxE* gene is expressed from the earliest stages of gonadogenesis, null mutant phenotypes are not apparent until the onset of metamorphosis [29].

In vertebrates, Group E genes are required in neural crest cells that contribute to the PNS [3, 61, 62] and we suggest the spider orthologue may have a similar function in the mechanoreceptors. These receptors are distributed all over the body, but the trichobothria only appear on the extremities of the limbs [63] where they differentiate from PNS progenitors.

Finally, the expression of *ptSoxF-2* is only detected at stage 9, in single dots at the tips of the L1 segment limb buds (Figure 8C). In *D. melanogaster* the *SoxF* gene is expressed in the embryonic PNS [56] and plays a role in the differentiation of sensory organ precursors [31], whereas in *A. mellifera* the *SoxF* orthologue is expressed ubiquitously throughout the embryo [13]. In *T. castaneum*, *E. kanangrensis* and *G. marginata* [45] *SoxF* expression is also associated with the embryonic limbs, again suggesting that this was an ancestral function of this Sox family in the Euarthropoda.

Taken together, our study expands our understanding of a highly conserved family of transcriptional regulators that appear to have played prominent roles in metazoan evolution. Our analysis indicates that the classification of Sox genes in the invertebrates appears to be robust and that genes in all Groups have aspects of their expression patterns that suggest evolutionary conservation across the Bilateria. In particular, it is becoming increasingly clear that a *SoxN* orthologue (SoxB1 in vertebrates) has a prominent role in the earliest aspects of CNS development. The finding that a *D/Sox21-b* class gene is implicated in the segmentation of both long and short germ band insects as well as the spider, and more widely in other arthropods [45], supports the view that formation of the segmented arthropod body plan is driven by an ancient mechanism [32], involving these Sox genes.

## Conclusions

Our analysis provides insights into the fate of duplicate genes in organisms that have undergone WGD. We find that virtually all the duplicates have been retained in the spider genome but the expression analysis suggests that some have been possibly been subject to subfunctionalisation and/or neofunctionalisation. It is interesting to note that in teleost fish, which have also undergone WGD events, the pattern we observe for the Sox family in spiders is mirrored, with considerable gene retention and lineage-specific neo-functionalisation [64]. Indeed, future functional studies in *P. tepidariorum* will help to reveal the precise roles played by Sox genes during spider embryogenesis and how this relates to other metazoans.

## Materials and Methods

### Genome analysis

TBLASTN searches of the *P. tepidariorum* and *S. mimosarum* genomes were performed with the HMG domain of mouse Sox2 (UniProtKB - P48432) at <u>http://bioinf.uni-greifswald.de/blast/parasteatoda/blast.php</u> and <u>http://metazoa.ensembl.org/Stegodyphus_mimosarum/Info/Index</u> respectively. Gene models were retrieved from the *P. tepidariorum* Web Apollo genome annotations via <u>https://www.hgsc.bcm.edu/arthropods/common-house-spider-genome-project</u> and from <u>http://metazoa.ensembl.org/Stegodyphus_mimosarum/Info/Index</u>. Sox gene sequences for other insects and vertebrates were retrieved from UniProt.

Multiple sequence alignments and phylogenetic analysis were performed with Clustal Omega [65] at <u>http://www.ebi.ac.uk/Tools/msa/clustalo/</u> or with MUSCLE and PhyLM 3.0 [66, 67] at <u>http://www.phylogeny.fr/index.cgi</u>. Pairwise sequence alignments were performed with SIM [68] at <u>http://web.expasy.org/sim/</u>.

### Synteny analysis of Sox genes in *P. tepidariorum*

The synteny of Sox genes was analysed to determine whether Sox genes were duplicated in the proposed WGD [41]. AUGUSTUS gene models in *P. tepidariorum* are already mapped against the DoveTail/HiRise genome assembly [41] and using these data the locations of Sox genes along with five upstream and five downstream flanking genes were compared. Gene models were removed if they were partial, chimeric or artefacts of the AUGUSTUS annotation to the HiRise assembly. To infer putative homology of flanking genes, their protein sequences were compared with BLASTP to the NCBI non-redundant protein sequence database [69].

### Embryo collection and procedures

Embryos were collected from adult female spiders from our temperature controlled (25°C) laboratory culture at Oxford Brookes University. Embryos at stages 5 to 12 were fixed as described in [70] and staged according to [71].

### In situ hybridisation

RNA in situ hybridisation was carried out as in [70], with slight modifications. Proteinase K treatment and post-fixations steps were omitted, the probes were heated to 80°C for 5 minutes and immediately put on ice before adding to the pre-hybridization buffer. Nuclear staining was performed by incubation of embryos in 1 μg/ml 4-6-diamidino-2-phenylindol (DAPI) in PBS with 0.1% Tween-20 for 15 minutes. Embryos were mounted in glycerol on Poly-L-lysine (Sigma-Aldrich) coated coverslips, where the germband tissue attaches making it easier to remove the yolk before imaging. Images were taken with an AxioZoom V16 stereomicroscope (Zeiss) equipped with an Axiocam 506 mono and colour digital camera. Brightness and intensity of the pictures were adjusted in Corel PhotoPaint X5 (CorelDraw).

### Gene isolation and cloning

Gene-specific cDNA fragments were amplified with primers designed with Primer Blast (https://www.ncbi.nlm.nih.gov/tools/primer-blast/) and PCR products cloned in the pCR4-TOPO vector (Invitrogen, Life Technologies). The primers to generate probe fragments for RNA *in situ* hybridization were designed to regions outside the consensus HMG domain to produce DNA fragments between 500-800 bp. The probes were *in vitro* transcribed as described in [70]. Primers and fragments size are described in Supplementary Table 3.

## Supplementary Tables

**Supplementary Table 1. HMG-domain and, where available full-length protein sequences from *D. melanogaster, P. tepidariorum, S. mimosarum* and *M. musculus***. Gene indicates the proposed names (or defined names for mouse). DB_Name indicates gene or gene model name form databases. DB_ID is the gene or protein accession. Scaffold indicates chromosome or genomic scaffold location. Annotation is the designation from spider annotations.

**Supplementary Table 2.** Gene and scaffold IDs of Sox and linked genes in the *P. tepidariorum* genome.

**Supplementary Table 3.** Genes, primers sequences and sizes for all the fragments used for in situ hybridisations.

## End Matter

**Availability of data and material**

Gene models for *P. tepidariorum* and *S. mimosarum* were retrieved from <u>https://www.hgsc.bcm.edu/arthropods/common-house-spider-genome-project</u> and from <u>http://metazoa.ensembl.org/Stegodyphus_mimosarum/Info/Index</u>. Sox gene sequences for animals were retrieved from UniProt. The annotated *P. tepidariorum* genome is available at <u>https://i5k.nal.usda.gov/JBrowse-partep</u> and the assembly is deposited at NCBI: BioProject PRJNA167405 (Accession: AOMJ00000000).

## Funding

This research was funded by a CNPq scholarship to CLBP (234586/2014-1), a grant from The Leverhulme Trust (RPG-2016-234) to APM and AS, and in part by a BBSRC grant (BB/N007069/1) to SR.

## Authors' contributions

Experiments were performed by CLBP, SR, AS and DL. All authors contributed to data analysis and writing the manuscript.

## Acknowledgements

We thank Evelyn Schwager for assistance in identifying Sox genes in the *P. tepidariorum* genome. CLBP is immensely grateful for the help and discussion with the members of the Embryology course (Woods Hole – 2016), especially Joaquin Navajas Acedo (Stowers Institute – USA) for pushing to keep the ball rolling.

---

**Supplementary Figure 1.**
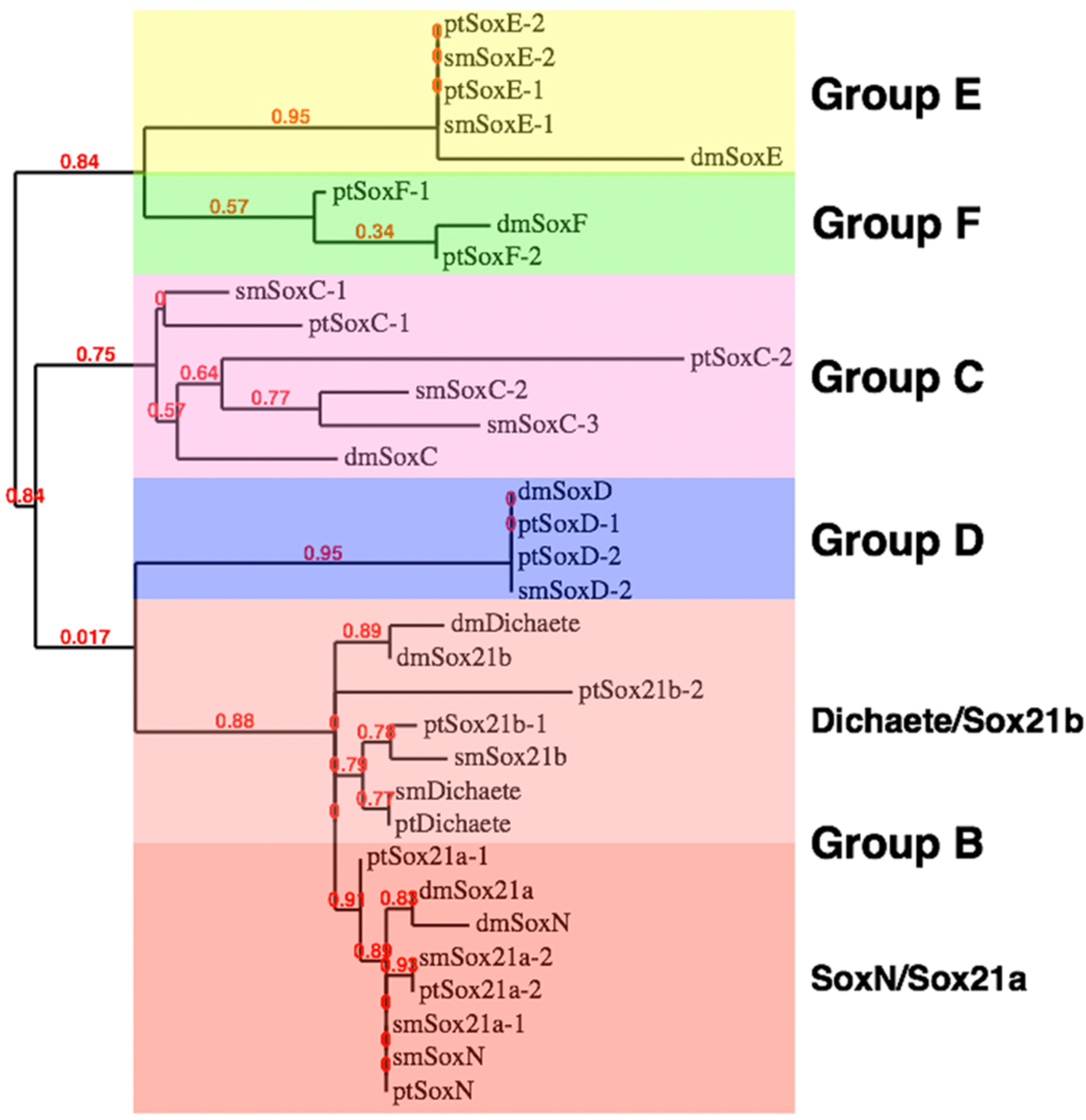
Phylogeny of Group B Sox HMG domains. PhyLM tree and multiple sequence alignment of group B HMG domains from *Mus musculus* (Mm), *Drosophila melanogaster* (Dm), *Anopheles gambiae* (Ag), *Tribolium castaneum* (Tc) *Parasteatoda tepidariorum* (Pt) and *Stegodyphus mimosarum* (Sm). Branch support values from PhyML are indicated in red. Arrow indicates the conserved Isoleucine reside indicative of invertebrate Dichaete/Sox21b class genes [15].

**Supplementary Figure 2.**
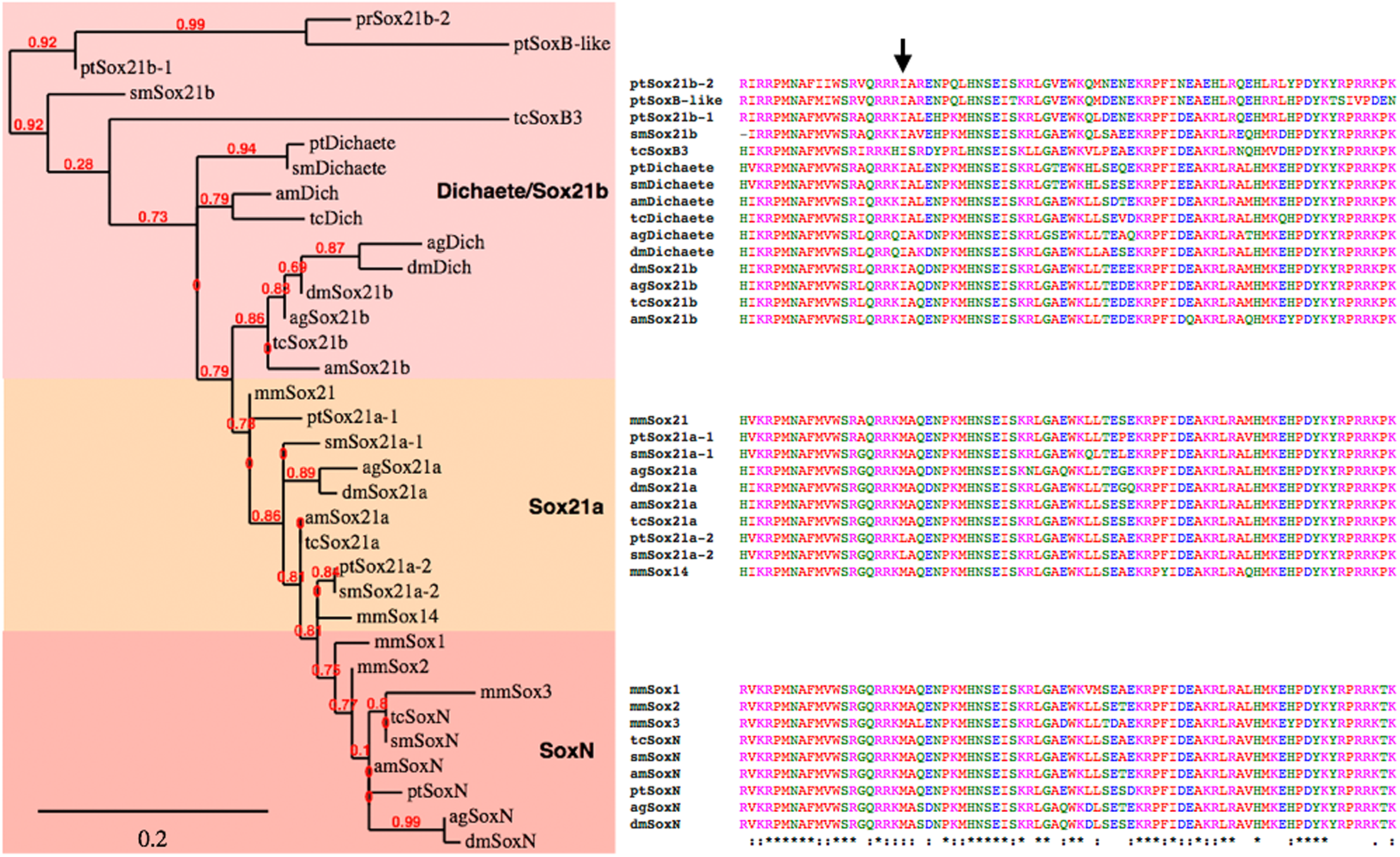
Phylogeny of full length Sox proteins from *Drosophila* and spiders. PhyLM tree of Sox genes from *D. melanogaster* (Dm), *P. tepidariorum* (Pt) and *S. mimosarum* (Sm) based on available full length protein sequence (Supplementary Table 1). Branch support values from PhyML are indicated in red.

